# Transcriptional dynamics of methane-cycling microbiomes are linked to seasonal CH_4_ fluxes in two hydromorphic and organic-rich grassland soils

**DOI:** 10.1101/2021.09.24.461182

**Authors:** Jana Täumer, Sven Marhan, Verena Groß, Corinna Jensen, Andreas W. Kuss, Steffen Kolb, Tim Urich

**Affiliations:** Institute of Microbiology, Center for Functional Genomics of Microbes, University of Greifswald, Greifswald, Germany; Institute of Soil Science and Land Evaluation, Soil Biology Department, University of Hohenheim, Stuttgart, Germany; Human Molecular Genetics Group, Department of Functional Genomics, University Medicine Greifswald, Greifswald, Germany; RA Landscape Functioning, Leibniz Centre for Agricultural Landscape Research (ZALF), Müncheberg, Germany; Thaer Institute, Faculty of Life Sciences, Humboldt University of Berlin, Berlin, Germany

## Abstract

Soil CH_4_ fluxes are driven by CH_4_-producing and -consuming microorganisms that determine whether soils are sources or sinks of this potent greenhouse gas. Using quantitative metatranscriptomics, we linked CH_4_-cycling microbiomes to net surface CH_4_ fluxes throughout a year in two drained peatland soils differing in grassland land-use intensity and physicochemical properties. CH_4_ fluxes were highly dynamic; both soils were net CH_4_ sources in autumn and winter and sinks in spring and summer. Despite similar net CH_4_ emissions, methanogen and methanotroph loads, as determined by small subunit rRNA transcripts per gram soil, differed strongly between sites. In contrast, mRNA transcript abundances were similar in both soils and correlated well with CH_4_ fluxes. The methane monooxygenase to methanogenesis mRNA ratio was higher in spring and summer, when the soils were net CH_4_ sinks. CH_4_ uptake was linked to an increased proportion of USCα and γ and pmoA2 *pmoA* transcripts. We assume that methanogen transcript abundance may be useful to approximate changes in net surface CH_4_ emissions from drained peat soils; high methanotroph to methanogen ratios would indicate CH_4_ sink properties. Our study shows the strength of quantitative metatranscriptomics; mRNA transcript abundance holds promising indicator to link soil microbiome functions to ecosystem-level processes.

## Introduction

CH_4_ is the second most important greenhouse gas, its global warming potential is 34 times higher than CO_2_ [1]. More than two-thirds of global CH_4_ emissions derive from microbial production in anoxic environments, such as wetland soils [2]. In past centuries, wetlands were commonly drained to gain agricultural land [3]. This lowers the water table, however, altering the physical properties of soils such as water content and oxygen availability. These altered soil properties, in turn, substantially affect the soil microbiota and activity and thus the soils’ greenhouse gas fluxes [4].

CH_4_-producing microbes (i.e., methanogens) are mostly anaerobic archaea that inhabit anoxic environments, e.g., wet rice paddies and wetland soils [2, 5]. Four types of methanogens can be characterized according to their substrate specificity. Acetoclastic methanogens utilize acetate, hydrogenotrophic methanogens utilize H_2_/CO_2_ and formate, and methylotrophic methanogens utilize methanol/methylamines to form CH_4_ [5]. Recently, methoxydotrophic methanogens that utilize methoxylated aromatic compounds were proposed as a novel methanogenic group [6, 7]. In wetlands, acetoclastic and hydrogenotrophic methanogens are considered the predominant sources of CH_4_ [5, 8]. However, recent research indicates that methanogenesis from methylated compounds also contributes to CH_4_ emissions from wetlands [9, 10].

Up to 90% of CH_4_ produced in oxygen-limited soils can be mitigated through oxidation by aerobic methanotrophs [11]. These methanotrophs are bacteria within the lineages *Alphaproteobacteria, Gammaproteobacteria*, and *Verrucomicrobia* [12]. CH_4_ oxidation can also be conducted anaerobically by bacteria in the NC10 phylum and archaea in the ANME group. Anaerobic oxidation of CH_4_ couples the reduction of other electron acceptors such as nitrite (NC10) or nitrate (ANME-2d) or ferric iron to the oxidation of CH_4_ [13–15]. Aerobic methanotrophs are considered the main oxidizers in wetland soils since alternative electron acceptors favoring anaerobic methanotrophs are often scarce in wetland soils. Besides their CH_4_ filter function, methanotrophs provide the only known biological sink for atmospheric CH_4_ [16]. However, it is not fully understood which microorganisms oxidize CH_4_ at atmospheric concentrations in soils. Bacteria of upland soil clusters (USC-α) and USC-γ have been identified as likely important atmospheric CH_4_ oxidizers in upland soils [12, 17, 18], while well-known methanotrophic lineages also account for CH_4_ oxidation in anoxic paddy soils [19].

Methanogens and aerobic methanotrophs differ substantially in their oxygen and substrate preferences. Generally, wet and anoxic conditions favor methanogenesis, whereas dry and oxic conditions favor aerobic oxidation of CH_4_. Drained peatland grasslands are highly dynamic across seasons in terms of water content and oxygen availability. Presumably, the interaction of methanogens and methanotrophs determines whether a wetland soil acts as a source or a sink for CH_4_ [20]. Linking CH_4_-cycling microbiome dynamics to CH_4_ fluxes, especially at the transcriptional level, has seldom been achieved [21].

DNA- and RNA-based meta-omics techniques have provided insight into the microbiome compositions of soils. However, DNA is stable over the long-term; extracted soil DNA may therefore partially originate from persistent extracellular DNA of dead organisms [22, 23]. In contrast, ribosomal RNA (rRNA) acts as a proxy for ribosomes and can provide a more accurate picture of the active microorganism community composition in soils. Still, the validity of rRNA as an indicator for the microbial activity of soils is questionable. For instance, dormant cells can contain high loads of ribosomes [24, 25]. Hence, although rRNA is a good proxy for potential soil microbiome composition, it may not relate well to microbial activity and ecosystem processes. The simultaneous sequencing of mRNA and rRNA can overcome this issue [26] because messenger RNA (mRNA), can serve as a proxy for potential transcriptional activity. Other metatranscriptomic studies indicate that mRNA is more responsive to environmental factors than rRNA [27, 28]. We thus aim to explore differences between small subunit (SSU) rRNA and mRNA transcripts of the CH_4_-cycling microbiomes and their links to CH_4_ fluxes. Furthermore, utilizing double RNA metatranscriptomics, it is possible to assign sequences to all three domains of life (bacteria, archaea, and eukaryotes) and to simultaneously assess their transcriptional activity [26].

Another drawback of meta-omics techniques is that they yield only relative abundances. However, the relationship between absolute abundances and relative abundances is not predictable [29]. It is thus hard to relate ecosystem processes to relative abundances. Studies have applied absolute quantification for metatranscriptomes in marine microbiomes [30, 31]. Recently, a quantification approach that uses total RNA to infer absolute from relative abundance was developed for metatranscriptomics [32]. This approach uses total RNA to infer absolute from relative abundance.

In this study, we aimed to link transcriptional dynamics of methane-cycling microbiomes to CH_4_ fluxes in two drained peatland soils. We used quantitative metatranscriptomics to analyze ribosomal rRNA and mRNA [26, 32] of 60 soil samples from drained peatland taken from different soil depths during autumn, winter, spring and summer. In addition, we measured CH_4_ and CO_2_ net surface fluxes from the two sites. We aimed to (a) evaluate the RNA content of the soils as a marker for microbial activity, (b) examine the CH_4_ fluxes of the two soils throughout a year, (c) study the composition and abundance of rRNA SSU and mRNA transcripts of CH_4_-cycling microbes, (d) link microbiome composition of CH_4_-cycling organisms to CH_4_ fluxes in drained peatland soils across seasons.

## Materials and Methods

### Site description

The experiment was conducted in the framework of the Biodiversity Exploratories project for long-term functional ecosystem research [33]. Samples were taken at two grassland sites (LI and HI) located in the Biosphere Reserve „Schorfheide-Chorin”(Table S1). Both sites are drained peatlands with a histosolic soil type (according to WRB 2015 [34]). The upper 30 cm of the peat soils was highly degraded. The two sites differ in the intensity of grassland management; the low land-use intensity site (LI) was mowed once or twice a year, while the high land-use intensity site (HI) was grazed by cows and additionally mowed sometimes once a year. Vegetation on LI was dominated by *Poa trivialis* (60%) and *Alopecurus pratensis* (25%); vegetation on HI was dominated by *Poa pratensis* aggr. (32 %), *Trifolium repens* (15%) and *Agrostis stolonifera* (10%).

### Soil Sampling

On each site, an area of 1 m x 7 m was sampled at all four seasons: autumn (11/09/2017), winter (03/08/2018), spring (05/30/2018), summer (09/13/2018). At each sampling date, three samples were taken between 12:00 and 13:00 at each site from the upper 10 cm and from the 20-30 cm layer. Each sample was a composite of soil cores from three locations. In spring, additional samples were taken at sunrise (05:00) and sunset (21:30). Samples for RNA, ammonium (NH_4_^+^), and nitrate (NO_3_^−^) extraction were immediately frozen at −80°C and subsequently stored as follows: RNA: −80°C, NH_4_^+^, and NO_3_^−^ −20°C. Samples for determination of C_mic_, N_mic_, pH, and soil water content were transported on ice and subsequently stored at −20°C. Redox potentials were measured with Mansfeld redox electrodes with an Ag/AgCl-reference electrode and a handheld ORP-meter GMH3531 (ecoTech, Bonn, Germany). For equilibration. the electrodes were placed in the soil 24 hours before sampling. Redox potentials were measured at soil depths of 5 cm and 25 cm.

### Determination of soil properties

Gravimetric soil water content was determined by drying 3–6 g soil at 65 °C to constant weight. Soil pH was determined by mixing 10 g dried sieved soil with 25 ml 0.01 M CaCl_2_ solution; pH of the suspension was then measured with a glass electrode (pH Electrode LE438, Mettler Toledo, Columbus, OH, USA). For total carbon and total nitrogen, samples were sieved (< 2 mm) and air-dried, ground in a ball mill (RETSCH MM200, Retsch, Haan, Germany), and analyzed in an elemental analyzer (VarioMax, Hanau, Germany) at 1100 °C. Inorganic carbon was determined with the same elemental analyzer after the organic carbon had been removed by combustion of soil samples at 450 °C for 16 h. Organic carbon concentration was calculated as the difference between total carbon and inorganic carbon. Microbial biomass carbon (C_mic_) and nitrogen (N_mic_) were determined by the chloroform-fumigation-extraction method (CFE), [35]. For this, frozen soils were thawed (at 4°C for 10 h), then 5 g field moist soils were fumigated with ethanol-free CHCl_3_ for 24 hours in a desiccator. C and N were extracted with 40 ml 0.5 M K_2_SO_4_, shaken horizontally (30 Min, 150 rpm), and centrifuged (30 Min, 4400 g) to separate extract from the soil. Non-fumigated soil samples were treated identically. Aliquots of the extracts were dissolved (1:4 extract:deionized. H_2_O) and measured on a TOC/TN analyzer (Multi N/C 2100S, Analytik Jena AG, Jena, Germany). A kEC factor [36]and a kEN factor [37] were used to calculate C_mic_ and N_mic_, respectively. Extractable organic carbon (EOC) and extractable nitrogen (EN) concentrations were determined on the non-fumigated samples. Mineral nitrogen in the forms of ammonium (NH_4_^+^) and nitrate (NO_3_^−^) was determined in the non-fumigated, non-diluted extracts with an AutoAnalyzer 3 (Bran & Luebbe, Norderstedt, Germany).

### Gas Fluxes

On each sampling date, gas emissions were measured 15 – 24 times throughout the day using the closed chamber method. Excessive vegetation was removed before pressing the stainless steel chambers (A = 150 cm^2^, V = 1800 ml) into the soil [38]. Four chambers per sampling per site were used. Gas samples (12 ml) were taken with syringes from the headspace immediately, 20, 40, and 60 minutes after closing the chambers via a three-way stopcock, and transferred into pre-evacuated exetainers (5.9 ml, Labco Lt, UK). Gas concentrations were measured on an Agilent 7890 gas chromatograph equipped with a flame ionization detector (for CH_4_) coupled with a methanizer (for CO_2_) (Agilent Technologies Inc., Santa Clara, CA, USA). Gas flux rates were calculated by the slope of the regression line of a linear regression of the gas concentration against time [17].

### RNA extraction, library preparation, and sequencing

Total nucleic acids were extracted using a phenol/chloroform/isoamylalcohol protocol [26]. The extracts were subsequently treated with DNase to remove DNA (DNase I, Zymo Research, Freiburg, Germany). RNA concentrations were measured with the Qubit® RNA HS Assay Kit (Qubit®3.0 Fluorometer, Invitrogen, Waltham, MA, USA.). RNA extracts were cleaned with the MEGAclear kit (Thermo Fisher Scientific, Waltham, MA, USA); quality of the RNA was verified by agarose gel electrophoresis and bioanalyzer (2100 Bioanalyzer, Agilent, Santa Clara CA, USA). We enriched the mRNA fraction and diluted inhibitory substances in the RNA extracts using the MessageAmp™ II-Bacteria RNA Amplification Kit (Thermo Fisher Scientific, MA, USA, input: 12.5 ng RNA). This method was previously validated for the preparation of metatranscriptomes [39]. Sequencing libraries were prepared with NEBNext® Ultra™ II RNA Library Prep Kit for Illumina® (New England Biolabs, Ipswich, MA, USA; input 60 ng). Manufacturer’s instructions were followed except for Step 4, where fragmentation time was adjusted to 3 min and a size selection step with HighPrep™ PCR beads (MagBio Genomics Inc., Gaithersburg, USA) was introduced (desired insert size 250 bp). Libraries were paired-end sequenced with an Illumina Next Seq 550 System using the NextSeq 500/550 High Output Kit v2.5 (300 Cycles) (Illumina, San Diego, CA, USA).

### Bioinformatic processing and Statistics

Reverse and forward sequences were overlapped with a minimum overlap of 10 or 5 bp with FLASH [40]. The sequences were filtered to a minimum mean quality score of 25 with PrinseqLite [41]. Sequences were then sorted into SSU, LSU, and non-rRNA fractions with SortMeRNA [42]. The SSU fraction was randomly subsampled to 200000 sequences with USEARCH [43]. Sequences were taxonomically classified against the SilvaMod128 databases [44] with BlastN [45] using a lowest common ancestor (LCA) algorithm in MEGAN (min score 155; top percent 2.0; min support 1, [46]). The non-rRNA fraction was aligned against the NC_nr database (retrieved 12/03/2020) with Diamond [47]. The sequences were taxonomically and functionally aligned with LCA in MEGAN (2011, min score 155; top percent 4; min support 1, [46]). Absolute abundances were calculated from read counts according to Söllinger *et al*. [32]. At the mRNA level, methanogenesis transcripts refer to sequences assigned to the SEED category “methanogenesis” (with subcategory “methanogenesis from methylated compounds”). Methanotrophy transcripts refer to sequences assigned to the SEED category “Particulate methane monooxygenase (pMMO)”. To classify *pmoA* sequences, the non-rRNA fraction was blasted against a *pmoA* database [48] and taxonomically classified with MEGAN as described in reference [48].

Statistical analyses were performed in R [49]. Distance based redundancy analysis was performed on the Bray Curtis dissimilarity matrix read counts of the 60 samples (function “dbrda” in the vegan package). Counts were Hellinger-transformed beforehand. We tested the following parameters: site (HI; LI), depth (“0-10 cm”, “20-30 cm”), season (“autumn”, “winter”, “spring”, “summer”), temperature, water content, nitrite, and nitrate. Continuous variables were z-scaled. Statistically significant categories between seasons and depths (within sites) were assessed by ANOVA and subsequent post-hoc Tukey test or multiple comparison test after Kruskal-Wallis at p < 0.05 level.

## Results and discussion

### Net surface gas fluxes have similar patterns across the year at both grassland sites

CH_4_ and CO_2_ fluxes were highly dynamic across the year (Figure 1). While the soils emitted CH_4_ in autumn and winter, they took up CH_4_ in spring and summer (Figure 1a). CO_2_ fluxes showed an opposite trend, with higher CO_2_ emissions in spring and summer than in autumn and winter (Figure 1b). The opposing trends of CO_2_ and CH_4_ fluxes reflected the changes in soil physicochemical properties across the year (Table S2). High water content and low redox potentials in autumn and winter favored anaerobic microbial processes, such as methanogenesis, while at the same time hampering aerobic microbial processes such as respiration. In contrast, soils had lower water content and positive redox potential in spring and summer, favoring aerobic respiration over anaerobic degradation processes. Generally, mean CO_2_ net surface emissions were about 1.5 times higher than IPCC default emission factors [50, 51]. Relatively high CO_2_ emissions have also been reported in other drained, nutrient-rich grasslands in Germany [52]. Our observed higher emissions may have been due to the highly degraded peat at the studied site. Soils with highly disturbed peat have been reported to have higher CO_2_ emissions than less degraded peat soils [52].

**Figure 1.**
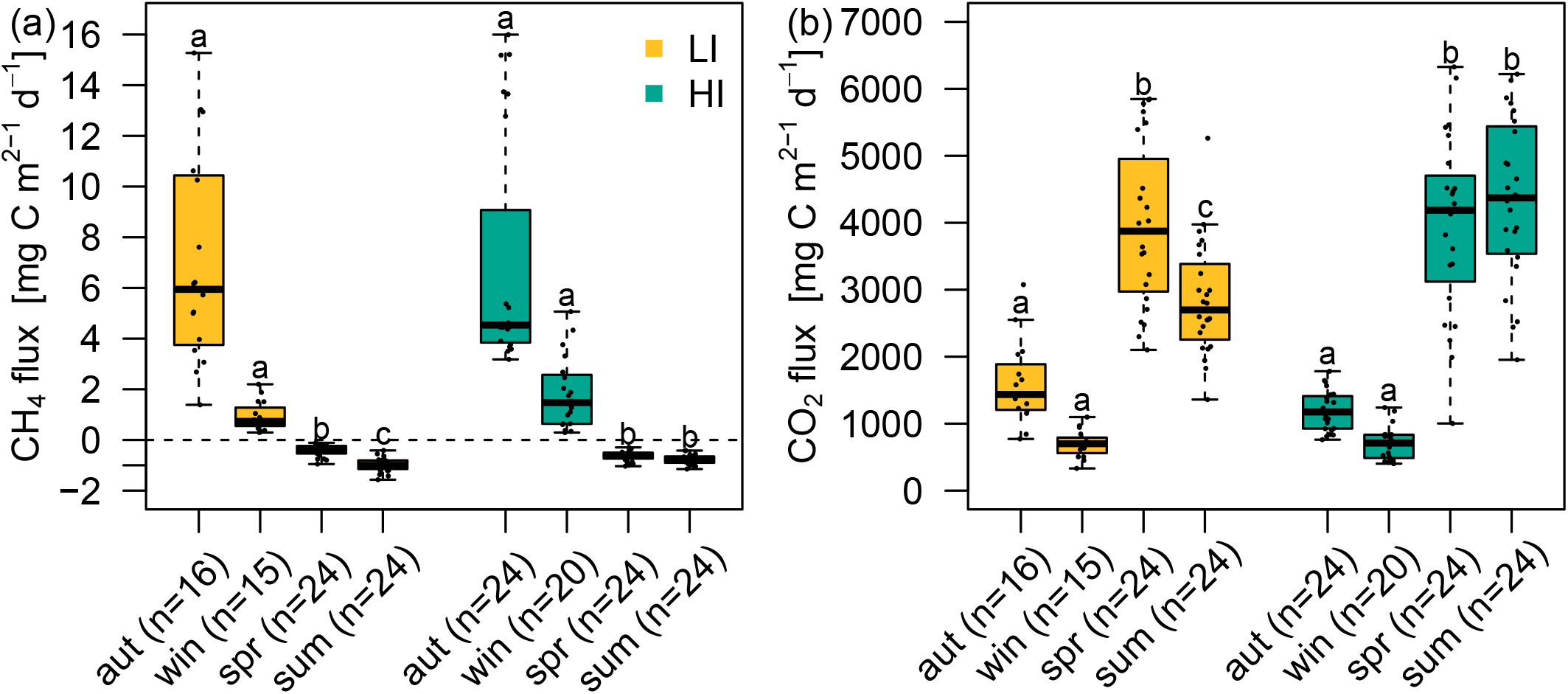
Net surface gas fluxes over the year. Gas fluxes of CH_4_(a) and CO_2_ (b) in mg C m^2^ d^−1^ at the grassland site with low (yellow, LI) and high (turquoise, HI) land-use intensity in autumn (aut), winter (win), spring (spr), and summer (sum). Boxes show the 25th and 75th percentiles and the lines inside the boxes indicate the median. Dots denote the individual measurements. Statistically significant categories (denoted with letters a,b and c) between seasons (per site) were tested with multiple comparison test after Kruskal-Wallis at p < 0.05 level. LI = low land-use intensity site, HI = high land-use intensity site.

In contrast, net surface CH_4_ emissions rates in autumn and winter were lower compared to IPCC default emission factors [50, 51]. However, we measured emissions at only four sampling times and may have not accounted for high emissions after heavy rainfall events. Net CH_4_ uptake rates in spring and summer were in the range of other grasslands and temperate ecosystems [53] and even slightly higher than in pastures [54]. In 2018, weather in spring and summer was dry and hot compared to the long-term average, leading to low soil water content (Table S2), favoring CH_4_ oxidation. The soil water content of the upper layer was mostly within the optimal range for atmospheric CH_4_ oxidation [55].

Our results underscore the highly variable greenhouse gas emissions from temperate drained peatlands and their dependence on dynamic soil physicochemical properties, clearly linked to seasons. Moreover, such sites can be net sinks for CH_4_ as well as net sources. This versatility regarding CH_4_ and source functions needs to be accounted for when estimating global carbon budgets.

### Linking metatranscriptomics and microbial biomass

We quantified total RNA content in the soils as a proxy for microbial biomass (Figure 2a). As expected, in most seasons the topsoils contained more RNA than the subsoils, especially the LI soil. The RNA content varied between seasons and was highest in winter and autumn in the topsoil and subsoil, respectively. Generally, RNA content was positively correlated with water content (r = 0.52, p < 0.001). Total RNA per gram soil was positively correlated with microbial biomass carbon and nitrogen (r_Cmic_^2^ = 0.29, r _Nmic_^2^ = 0.46, p < 0.001, figure 2). Interestingly, RNA content correlated better with N_mic_, likely due to the high N content of the RNA. This finding supports the validity of RNA as a proxy for living microorganisms and enables the use of RNA content to infer absolute transcript abundances from relative abundances [32]. Through this quantitative approach, we overcame challenges typically associated with the interpretation of relative abundance in ‘meta-omics’ datasets. A recent study used this quantitative approach and found that absolute transcript abundance correlated better to processes than relative transcript frequencies [32].

**Figure 2.**
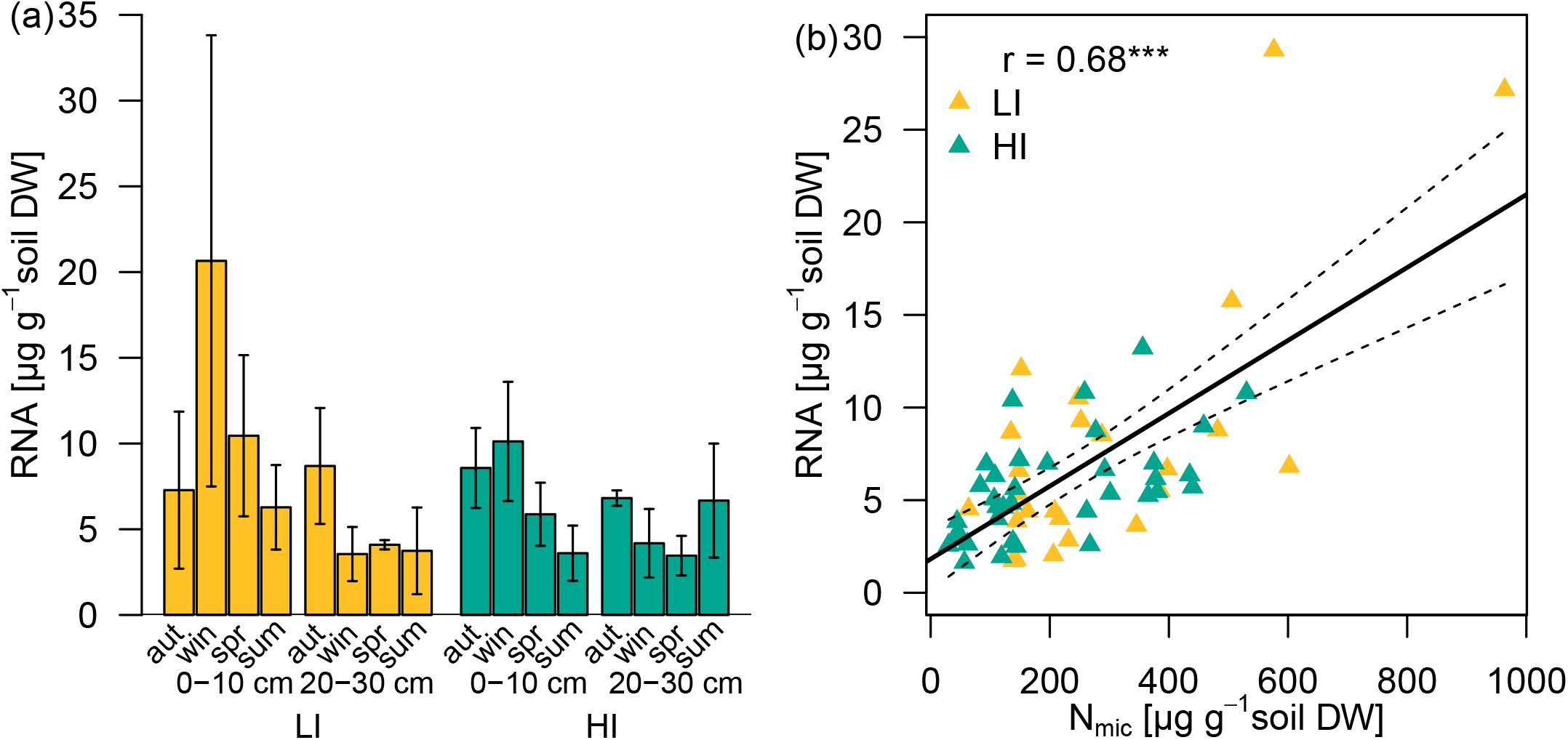
RNA and microbial nitrogen content. RNA content (μg g^−1^ soil DW) across seasons in the upper (0-10 cm) and the deeper soil layer (20-30 cm) at the site with low (yellow) and high (turquoise) land-use intensity (a), and correlation between RNA and microbial nitrogen content (N_mic_) per g soil (b). Linear regression RNA = 1.81824+ 0.01968 N_mic_, df = 58. LI = low land-use intensity site, HI = high land-use intensity site. R denotes the Pearson correlation coefficient. Significance codes: *** = p< 0.001, n= 60. Abbreviations: aut = autumn, win = winter, spr = spring, sum: summer, LI = low land-use intensity site, HI = high land-use intensity site, DW = dry weight.

### Spatio-temporal dynamics in methane-cycling (micro-)biome

The relative abundances (RAs) of SSU rRNA transcripts of the three domains ranged from 54.6% to 95.4% for Bacteria, 0.7% to 10.1% for archaea, and 1.7% to 41.3% for Eukaryotes (Table S3 and Table S4). The community composition of all taxa in the soil samples exhibited a clear site- and depth-specific pattern (Figure 3a). The tested parameters explained 52.7% of the variance in the community composition of all taxa (Table S2). Site and depth had the greatest explanatory power (20.0% and 19.6% respectively, p<0.001, Table S4). Site-specific differences could likely be attributed to site-specific soil properties, such as pH, texture, organic carbon, and nitrogen content (Table S1). Depth is generally considered to be associated with differences in oxygen and nutrient availability. Season accounted for 6.0% of the variation (p < 0.01). This effect likely resulted from varying precipitation, water table depth, and plant growth activity throughout the year.

**Figure 3.**
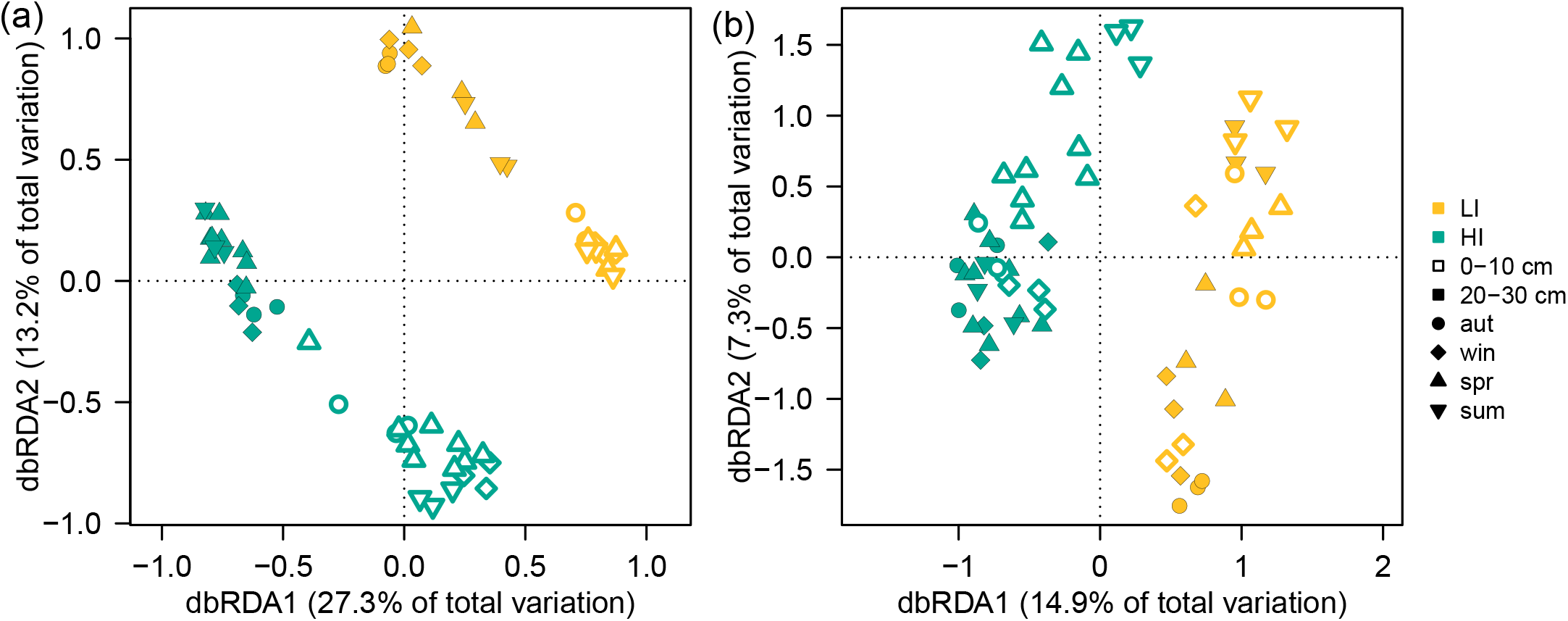
Soil (micro-)biome composition across sites and soil depths. Distance-based redundancy analysis (dbRDA) of the Bray-Curtis dissimilarity matrix of all 39854 bacterial, archaeal and eukaryotic taxa (a) and the 287 CH_4_-cycling archaea and bacteria (b) in the grassland sites with low (yellow) and high (turquoise) land-use intensity. Samples from the upper (0-10 cm) and the deeper soil layer (20-30 cm) are depicted with hollow and filled circles, respectively. Samples from autumn, winter, spring and summer are depicted as circles, diamonds, upward-pointing triangles and downward-pointing triangles, respectively. Abbreviations: aut = autumn, win = winter, spr = spring, sum = summer, LI = low land-use intensity site, HI = high land-use intensity site.

We further identified and quantified three different groups of CH_4_-cycling microorganisms in the SSU sequences: (i) methanogenic archaea (RA: 0.01% to 1.99%), (ii) aerobic methanotrophic bacteria (RA: 0.02% to 1.39%), and (iii) anaerobic methanotrophic archaea and bacteria (RA: 0% to 0.12%, Table S3 and Table S4). When considering only CH_4_-cycling taxa (total of 287 identified taxa), the tested parameters explained 36.1% of the variance in community composition (Table S5). Site had the most explanatory power (14.0%, p < 0.001), whereas depth, season, and water content accounted for 5.7% (p < 0.001), 6.5% (p < 0.001), and 5.3% (p < 0.001) of the variance, respectively. The water content of soils directly affects O_2_ availability, which, in turn, is a fundamental factor shaping CH_4_-cycling community composition. O_2_ diffuses more slowly in water-saturated soils, which thus have lower O_2_ availability [56].

### Methanogen community composition influences transcriptional activity

Methane formation is a complex process involving several functional guilds that metabolically interact closely with each other [20]. We integrated the total RNA content and metatranscriptomes to infer SSU rRNA and mRNA transcript abundances per gram soil [32]. Most methanogen families in the soils were class II methanogens, e.g., *Methanosarcinaceae, Methanosaetaceae* (now *Methanotrichaceae*; Figure 4). Generally, these families possess more antioxidant features than class I methanogens [57]. The predominance of class II methanogens reflected the dynamic water and redox status in the studied soils (Table S2). Despite comparable CH_4_ emissions at both sites, the abundance of methanogens was 4.6 times higher in the soils of high land use intensity (HI) compared with that of low land use intensity (LI) (Figure 4a,b, p < .001). Interestingly, methanogenesis-related transcript abundances were not statistically different between sites (Figure 5a), suggesting lower transcriptional activity of the methanogens in the low land-use intensity (LI) grassland soil. The incongruity between methanogen SSU rRNA abundances and CH_4_ fluxes at the SSU level may also be attributable to differences in taxonomic compositions of methanogens. The strictly acetoclastic *Methanosaetaceae* (*Methanothrix*) were about 30 times more abundant in HI than in LI (Figure 4a, p < 0.01), especially in the deeper soil layer. *Methanosaeta* have lower growth rates but can grow at lower acetate concentrations than the metabolically diverse *Methanosarcina* [58]. Lower growth rates may lead to lower transcriptional activity and lower CH_4_ emissions. *Methanosaeta* often outcompete other acetoclastic methanogens at low acetate concentrations [58, 59]. Thus, the higher proportion of *Methanosaetaceae* may have indicated lower acetate availability in HI. The lower acetate availability in HI could thus have resulted in observed lower CH_4_ emissions despite the higher methanogen abundances compared with LI. Interestingly, the share of hydrogenotrophic methanogens, such as *Methanocellaceae, Methanoregulaceae* and *Methanobacteriaceae*, was higher in LI than in HI. The energy yield of hydrogenotrophic methanogenesis is larger than that of acetoclastic methanogenesis, which has the lowest energy yield of all methanogenic metabolisms [5, 60]. The varying proportions of acetoclastic and hydrogenotrophic methanogens and lower substrate availabilities at the two sites may explain our observed differences between CH_4_ emissions and methanogen abundances at the SSU level. Our results suggest that not solely the abundances but also the composition of methanogens influence net surface CH_4_ fluxes of soils.

**Figure 4.**
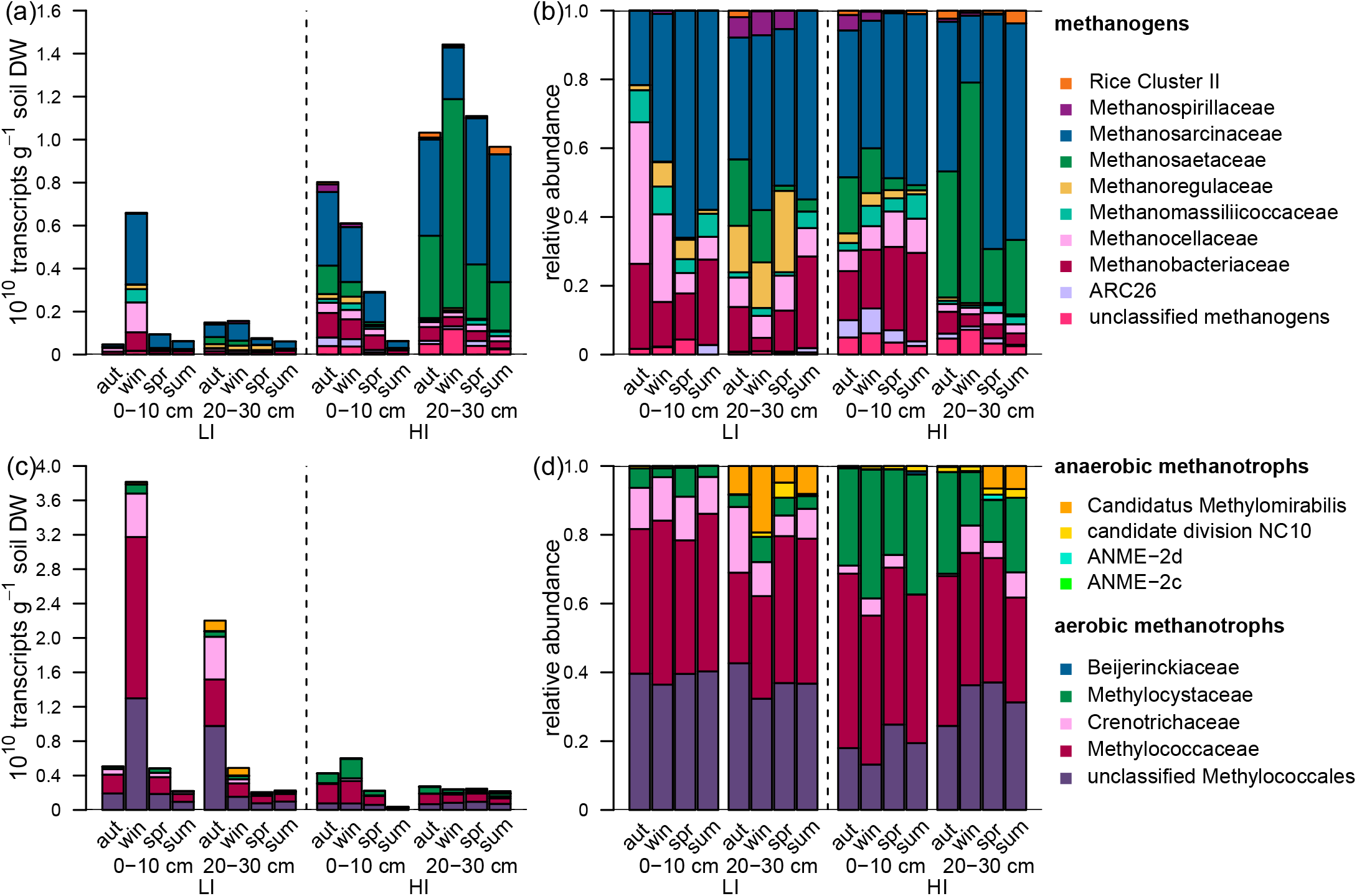
Absolute and relative abundances of CH_4_-cycling organisms across seasons and depths at small subunit (SSU) rRNA level. Absolute abundances (SSU transcripts g-1 soil DW) of methanogenic Archaea (a) and methanotrophic microorganisms (Archaea and Bacteria) (c) and the proportion of transcripts belonging to methanogenic Archaea (b) and methanotrophic microorganisms (d) normalized to the total amount of transcripts belonging to methanogenic Archaea and methanotrophic organisms. Columns show means across seasons of the upper (0-10 cm) and the deeper soil layer (20-30 cm) in the grassland site with low (LI) and high land-use intensity (HI). “unclassified methanogens” contains methanogens unclassified at the class level and low abundance methanogenic groups. “unclassified *Methylococcales*” contains *Methylococcales* unclassified at the family level and low abundance *Methylococcales* families. Bars represent the means of three replicates. Abbreviations: aut = autumn, win = winter, spr = spring, sum: summer, LI = low land-use intensity site, HI = high land-use intensity site, DW = dry weight. n= 60

**Figure 5.**
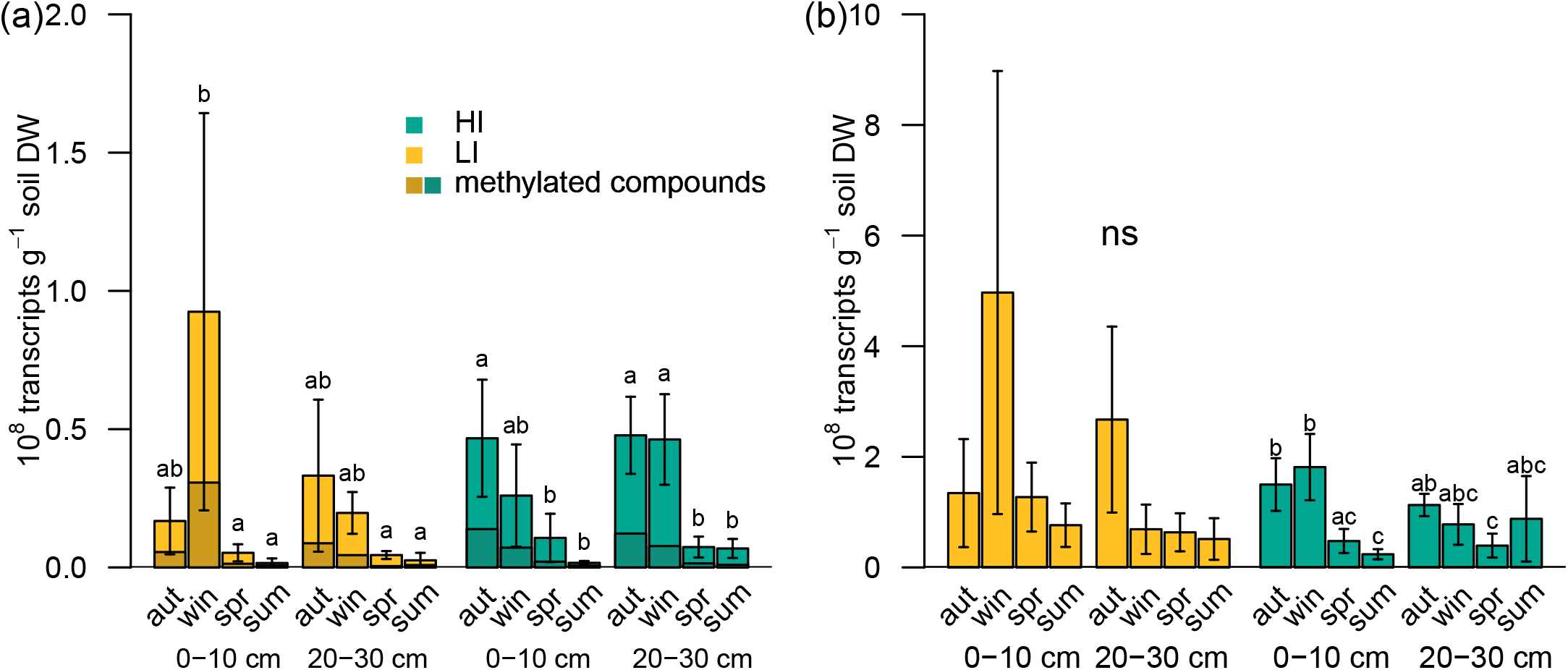
Abundances of CH_4_-cycling mRNA transcripts across seasons and depths. Transcript abundances (transcripts g^−1^ soil DW) of methanogenesis pathways (a) and pMMO (b) across seasons in the upper (0-10 cm) and the deeper soil layer (20-30 cm) the grassland sites with low and high land-use intensities. Columns and error bars represent the means and the standard deviations of three replicates, respectively. Statistically significant categories between seasons and depths (per site) were tested with ANOVA and subsequent post-hoc Tukey test at p < 0.05 level, ns = not significant, n= 60. Abbreviations: aut = autumn, win = winter, spr = spring, sum: summer, LI = low land-use intensity site, HI = high land-use intensity site, methylated compounds= methanogenesis from methylated compounds, DW = dry weight.

Methanogenesis from methylated compounds is the predominant source of CH_4_ in marine environments and its contribution to the global CH_4_ budget is considered to be small [5]. However, recent research suggests that it might be more important than previously assumed [9, 61–63]. The presence of *Methanomassiliicoccales* (up to 14% of the total methanogen community in a winter topsoil sample), and especially *Methanosarcinaceae*, points to methylated compounds as a substrate for methanogenesis at our sites (Figure 4a,b [62]). In support of this, the portion of mRNAs for methanogenesis from methylated compounds was up to 50% of all methanogenesis transcripts in some datasets (Figure 5a). Methanogenesis from methylated compounds in wetlands may thus be more important than previously thought [6, 9, 61–63].

### High methanotrophic diversity in the studied grassland soils

We detected aerobic and anaerobic methanotrophs belonging to *Methylococcaceae, Crenotrichaceae, Methylocystaceae* (aerobic), and NC10 and ANME (anaerobic) in all microbiomes (Figure 4c,d). Our studied sites had highly dynamic soil properties and fluctuating water levels throughout the year. Therefore, the soil provided versatile niches for different kinds of methanotrophs with distinct demands on CH_4_ and O_2_ levels. The majority of the detected methanotrophs belonged to the obligate methanotrophic *Methylococcaceae*. Normalized to all methanotrophic taxa, *Methylocystaceae* was more abundant in HI than in LI, while *Crenotrichaceae* were more abundant in LI (p < 0.05).

#### Anaerobic methanotrophs

Anaerobic methanotrophs were present mainly in the deeper soil layer, which was likely due to their sensitivity to oxygen [64]. Generally, there were more aerobic than anaerobic methanotrophs (Figure 4c,d). However, anaerobic methanotrophs accounted for up to 20% of all methanotrophs and consequently comprised a substantial part of the methanotroph community at the studied sites. Another study found a similar proportion of anaerobic methanotrophs (5.7-25.8%) in the upper 10 cm of urban wetlands [65]. Zhong *et al*. [66] found that the relative abundance of anaerobic methanotrophs in wetland soils was highest at the 50-60 cm depth. We investigated only to a depth of 30 cm. The proportion of NC10 with respect to all methanotrophs may have been even higher in the less oxygen-influenced deeper soil layer. However, redox potential alone cannot explain the distribution of NC10 in our study since abundances of NC10 showed divergent trends at the two sites. In HI, their abundance was highest in summer, despite rising redox potentials. This increase may have been due to the increase in soil nitrate content in spring and summer, probably a result of grazing and subsequent mineralization and N release from feces.

Predominant anaerobic methanotrophs were the nitrite-dependent NC10 bacteria, especially *Candidatus* “Methylomirabilis” (Figure 4c,d). They have also been identified as the most relevant anaerobic CH_4_ oxidizer group in freshwater marsh sediments [67]. Our results indicate *Ca*. Methylomirabilis is also the main anaerobic methanotroph in drained peatlands. We found archaeal CH_4_ oxidizers to be absent at most seasons and highest in abundance in spring in HI (Figure 4c,d), which coincides with a nitrate increase in this soil (Table S2). Another study found archaeal CH_4_ oxidizers more abundant than NC10 bacteria in a fertilized paddy soil [68]. Thus, high nitrate content or nitrogen input may favor archaeal over bacterial anaerobic CH_4_ oxidizers. Our results suggest that NC10 is a relevant methanotrophic group mitigating CH_4_ emissions from drained peatlands and that oxygen and substrate availability are important factors influencing their abundance.

#### Methanotrophs as CH_4_ filter and CH_4_ sink

Methanotrophs can oxidize CH_4_ at atmospheric and elevated CH_4_ concentrations [12]. In our study, mRNA methanogenesis transcripts correlated with PMO transcripts (r = 0.62, p < 0.05, Figure 6d). This relationship suggests that methanotrophs use CH_4_ derived from methanogenesis in the soil. They thus act as a filter mitigating CH_4_ emission to the atmosphere. In spring and summer, the studied sites were a CH_4_ sink, suggesting that the detected methanotrophs metabolized CH_4_ at atmospheric concentrations and below. Which organisms are responsible for atmospheric CH_4_ oxidation in soils remains unresolved since until recently atmospheric CH_4_ oxidizers resisted cultivation. However, atmospheric CH_4_ consumption in upland soils is assumed to be performed by bacteria of Upland soil clusters (USC) α and γ [12, 18]. The first isolated USCα methanotroph, *Methylocapsa gorgona*, can grow at both atmospheric and elevated CH_4_ concentrations [69]. Furthermore, *Methylocystis* strains can also oxidize atmospheric CH_4_ [70]. We did not detect an increase in known atmospheric CH_4_ oxidizers in the SSU transcripts. However, the proportion of mRNA transcripts belonging to USCα and γ increased in spring and summer (Figure 6). This supports the proposal that both groups are involved in atmospheric CH_4_ oxidation in temporarily wet grassland soils. Additionally, the relative abundance of *pmoA2* transcripts increased towards summer. *pmoA2* is part of the *pmoCAB2* operon, which encodes a pMMO capable of oxidizing CH_4_ at atmospheric levels [70]. Many type II methanotrophs contain *pmoA2* [71]. These groups could therefore also be responsible for atmospheric CH_4_ oxidation in the studied soils. The relative abundance of USCα and γ *pmoA* transcripts and *pmoA2* was up to 34%. Still, other type-I and type-II *pmoA* sequences dominated in the soils. Thus, also type I and type II methanotrophs may be involved in atmospheric CH_4_ oxidation. Recently, high-affinity CH_4_ oxidation in paddy soils was attributed to well-known CH_4_ oxidizers rather than USCα and γ [19]. In a former study, we found USCα and γ *pmoA* gene abundances to be explanatory factors for atmospheric CH_4_ oxidation in grassland soils of three different regions of Germany, including the Schorfheide region investigated in this study [17]. Despite the wide distribution of USCα and γ sequences, *pmoA* sequences detectable with A189f/mb661 primers were absent in most regions except for the Schorfheide grasslands. The A189f/mb661 primer pair amplifies a broad range of well-known *pmoA* sequences but discriminates USCα and γ sequences [72]. The presence of these well-known sequences in the Schorfheide was likely due to the hydromorphic character of the soils. This region contains many drained peatland soils with internal CH_4_ formation by methanogens. It is possible that different groups contribute to atmospheric CH_4_ oxidations in soils with internal CH_4_ formation (such as the drained peatlands investigated here), as compared to upland soils that are permanent net sinks of CH_4_. Possibly, USCα and γ methanotrophs are the predominant atmospheric CH_4_ oxidizers in such soils. In contrast, in soils with internal CH_4_ formation, even only in some seasons, other methanotrophs could also contribute to atmospheric CH_4_ oxidation, as suggested in [19].

**Figure 6.**
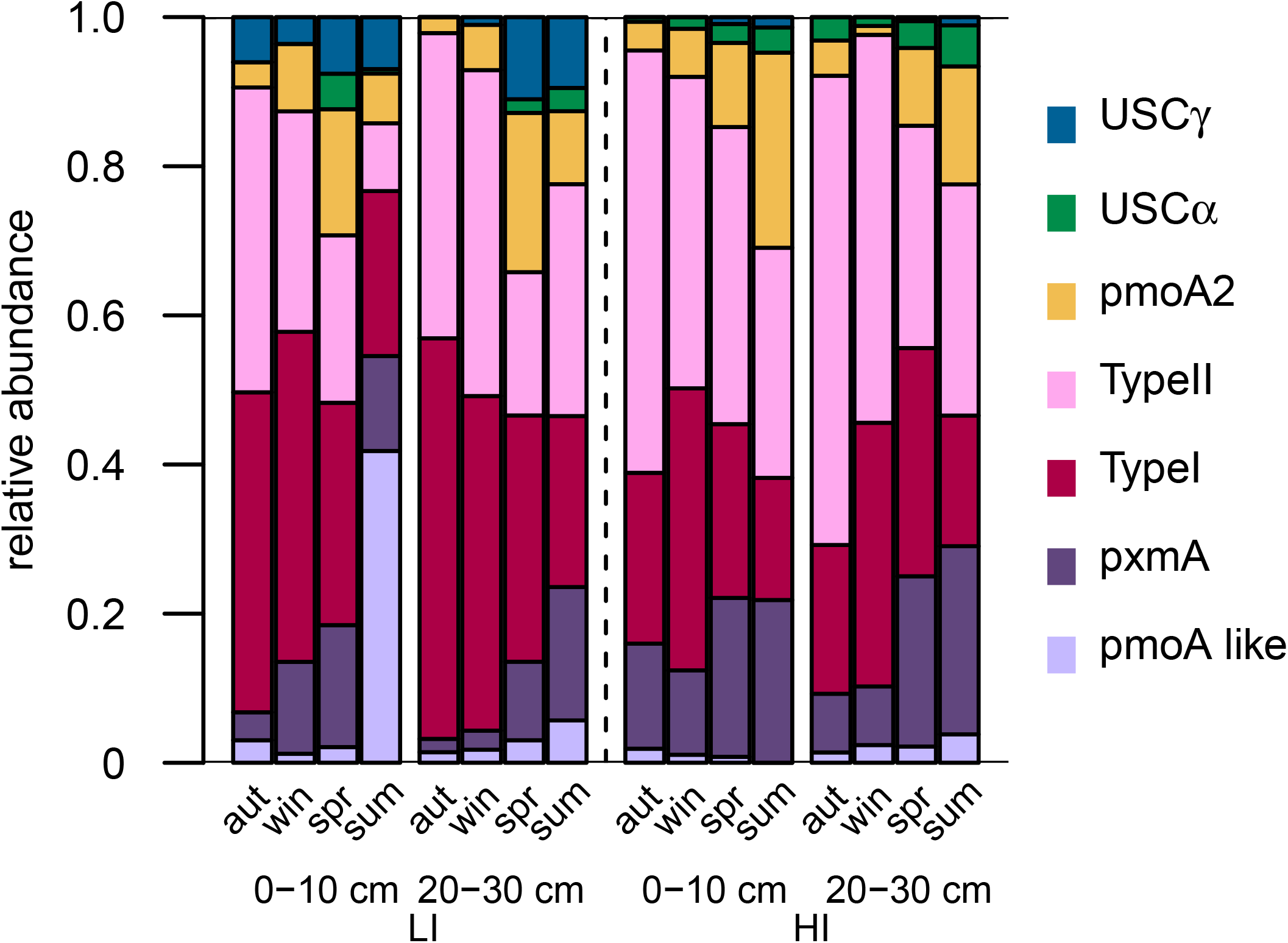
Composition of *pmoA* transcripts. Proportion of *pmoA* groups normalized to the total amount of *pmoA* transcripts. Columns show means across seasons of the upper (0-10 cm) and the deeper soil layer (20-30 cm) in the grassland sites with low (LI) and high land-use intensity (HI), n= 60. “pmoA_like” = unclassified *pmoA*-like sequences. Abbreviations: aut = autumn, win = winter, spr = spring, sum: summer, LI = low land-use intensity site, HI = high land-use intensity site.

### Functional transcript abundances as a proxy for soil net surface CH_4_ fluxes

Messenger RNAs of methanogenesis pathways were less abundant per gram soil in spring and summer than in autumn and winter at both sites and depths (Figure 5a). This drop in mRNA agrees with the cessation of CH_4_ emissions from the soils in spring in summer; both correlated significantly with each other (r = 0.82, p < 0.05, Figure 7b). In contrast, abundances of methanogen SSU transcripts and CH_4_ fluxes did not significantly correlate within either plot (Figure 7a). However, there was a site-specific trend. Our results indicate that quantified methanogenesis mRNA transcripts were better indicators of net CH_4_ fluxes than methanogen SSU rRNA transcripts. Our findings thus underscore studies that have found mRNA more responsive to environmental factors than rRNA [27, 28].

**Figure 7.**
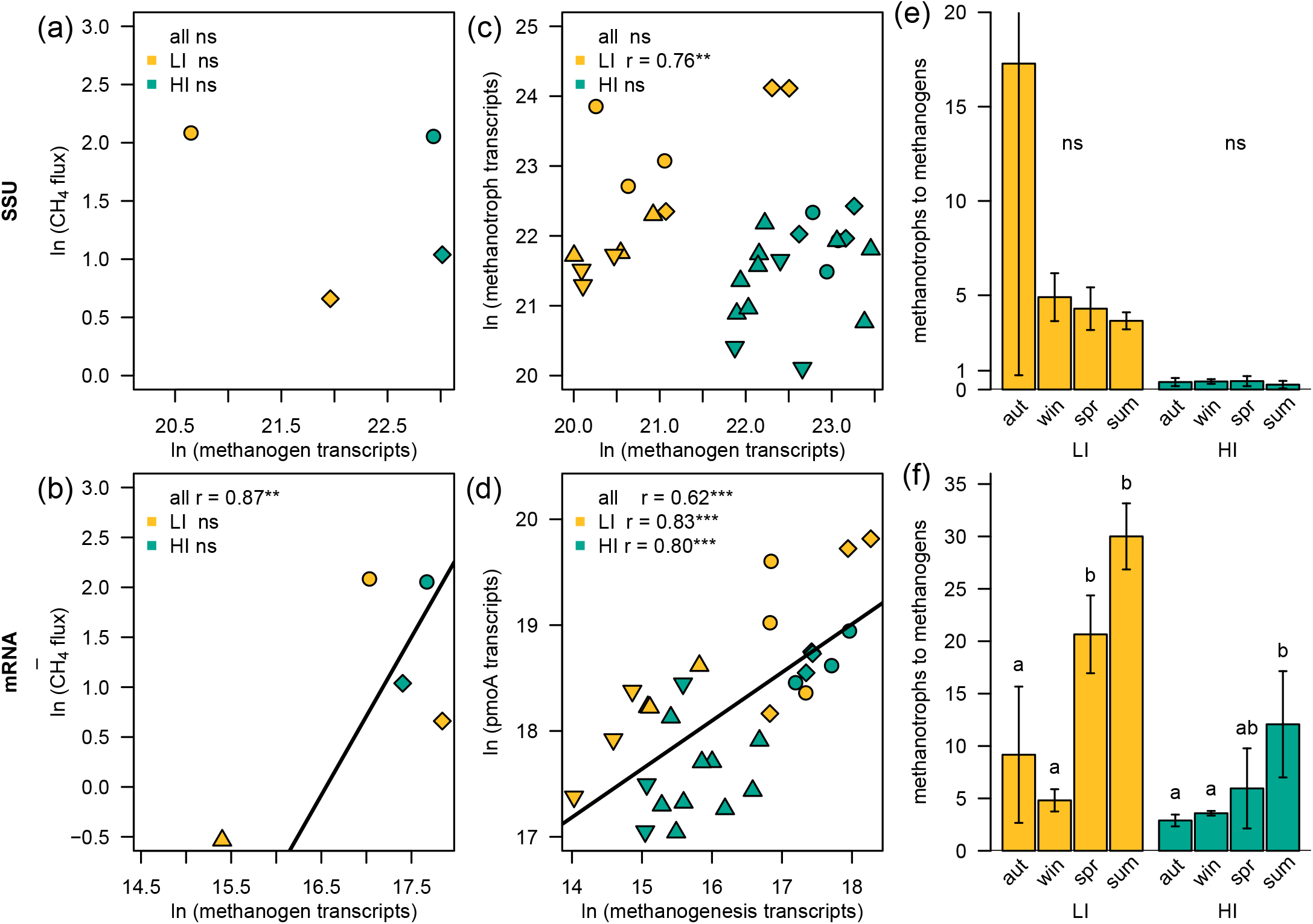
The link between methanogenesis gene transcripts, net surface CH_4_ fluxes, and methanotrophy gene transcripts. Linear correlation of absolute abundances of methanogen transcripts and CH_4_ flux at SSU level (a) and mRNA level (b). Linear correlation of absolute abundances of methanogen and methanotroph transcripts at SSU level (c) and mRNA level (d). Ratio of methanotroph to methanogen transcripts across the year at SSU rRNA level (e) and mRNA level (f) across seasons at the grassland sites with low and high land-use intensity. R denotes the Pearson correlation coefficient. Significance codes: * = p < 0.05, ** = p < 0.01, *** = p< 0.001, ns = not significant, n= 60. Statistically significant categories were tested with ANOVA and subsequent post-hoc Tukey test at p < 0.05 level. In (a) and (b) points represent seasonal means across both depths. In (c) and (d) points represent mean values per core (means of both depths). The different colors represent the site with low (yellow) and high (turquoise) land-use intensity. Samples from autumn, winter, spring and summer are depicted as circles, diamonds, upward-pointing triangles and downward-pointing triangles, respectively. Abbreviations: aut = autumn, win = winter, spr = spring, sum: summer, LI = low land-use intensity site, HI = high land-use intensity site.

Methanotrophs are important CH_4_ filters, mitigating soil CH_4_ emissions. The pMMO to methanogenesis mRNA ratio was higher in spring and summer than in autumn and winter (Figure 7f). However, the ratio of methanotroph to methanogen SSU transcripts did not permit conclusions about the soils’ CH_4_ fluxes (Figure 7e). The mRNA ratio may thus indicate whether soils are a CH_4_ source or sink. A high methanotroph to methanogen ratio indicates the soil serves as a CH_4_ sink, while a low ratio indicates the soil serves as a net CH_4_ source. Yet it is necessary to consider not only SSU rRNA transcripts but also mRNA transcripts. Furthermore, community composition could be an additional indicator, since a high proportion of atmospheric CH_4_ oxidizers were linked to net CH_4_ uptake of the soils.

Additionally, our results may suggest that abundance of methanogenesis related mRNAs is an estimator of soil CH_4_ fluxes. in soils with a low methanotroph to methanogen ratio. We used RNA sequencing, which is still restricted in terms of throughput. Two RT-qPCR studies found a relationship between *mcrA* transcript abundances and CH_4_ fluxes in a paddy soil and a peat bog [73, 74]. RT qPCR might thus be a tool to estimate CH_4_ fluxes from *mcrA* transcript abundances in combination with *pmoA* transcript abundances. However, large-scale studies are needed to further investigate the link between methanogen and methanotroph transcripts and CH_4_ fluxes across different soil types and seasons.

## Conclusions

CH_4_-cycling microbes interact with each other on a process level. These interactions among anaerobic archaea and aerobic and anaerobic bacteria determine the net surface CH_4_ fluxes from soils. This study is, to our knowledge, the first to use quantitative metatranscriptomics to link CH_4_ fluxes from grasslands with CH_4_-cycling microbiomes through all seasons of the year. We investigated two drained peatland soils with highly variable soil physicochemical properties, an associated dynamic microbial community composition, and net surface CH_4_ fluxes in both directions, at the same sites through the year. It is crucial to account for the observed ambiguous behavior of such drained wetlands when estimating the CH_4_ budgets of grassland ecosystems in regional and global surveys. We further validated that mRNA transcripts rather than SSU transcripts are necessary to link microbial activity to soil net CH_4_ fluxes. Particulate MMO transcripts were correlated to mRNA of methanogenesis pathways, indicating that methanotrophs acted as CH_4_ filters. Net CH_4_ uptake was linked to an increase in proportions of USCα and γ and *pmoA2* transcripts, supporting evidence that these groups are involved in atmospheric CH_4_ oxidation. Messenger RNA of methanogenesis pathways correlated with net CH_4_ fluxes. It may thus be feasible to estimate soil CH_4_ fluxes using *mcrA* transcript abundances, additionally considering *pmoA* transcripts. Our results suggest that the ratio of *pmoA* to *mcrA* transcripts is an easily applicable estimator of whether a soil acts as a sink or source of CH_4_. However, large-scale and long-term studies are needed to evaluate and to elaborate this topic further.

## Acknowledgements

We thank the managers of the three Exploratories, Miriam Teuscher, Juliane Vogt, Kirsten Reichel-Jung, Iris Steitz, Sandra Weithmann and all former managers for their work in maintaining the plot and project infrastructure; Christiane Fischer and Victoria Grießmeier for giving support through the central office, Andreas Ostrowski for managing the central data base, and Markus Fischer, Eduard Linsenmair, Dominik Hessenmöller, Daniel Prati, Ingo Schöning, François Buscot, Ernst-Detlef Schulze, Wolfgang W. Weisser and the late Elisabeth Kalko for their role in setting up the Biodiversity Exploratories project. We thank the administration of the Hainich national park, the UNESCO Biosphere Reserve Swabian Alb and the UNESCO Biosphere Reserve Schorfheide-Chorin as well as all land owners for the excellent collaboration. The work has been funded by the DFG Priority Program 1374 “Biodiversity-Exploratories” (DFG-KO2912/12-1, MA4436/2-1, UR198/3-1, UR198/5-1). Field work permits were issued by the responsible state environmental offices of Baden-Württemberg, Thüringen, and Brandenburg. We thank Markus Fischer and the botany core team for providing vegetation record data. We thank Lars Jensen and Lisa Hagenau for their support during library preparation and sequencing. We thank Juliette Blum and Fabian Stache (University Hohenheim) for performing microbial biomass analyses. We also are grateful for the support by ZALF technicians, especially Sigune Weinert and Paul Reim. We thank Uta Schumacher and Markus Rubenbauer for helping to coordinate the field work. And we thank Andrea Söllinger, Tilman Schmider, Sebastian Petters, Klary Rychly, Vjera Kovacevic, Micha Weil, Simon Weddell, Karolin Müller, Felix Müller and Eva Aderjan for help and support during field sampling. We also thank Mathilde Borg Dahl for advice with statistical analysis.

